# Investigating Brain Alterations in the Dp1Tyb Mouse Model of Down Syndrome

**DOI:** 10.1101/2023.07.26.550698

**Authors:** Maria Elisa Serrano, Eugene Kim, Bernard Siow, Da Ma, Loreto Rojo, Camilla Simmons, Darryl Hayward, Dorota Gibbins, Nisha Singh, Andre Strydom, Elizabeth M. C. Fisher, Victor L. J. Tybulewicz, Diana Cash

## Abstract

Down syndrome (DS) is one of the most common birth defects and the most prevalent genetic form of intellectual disability. DS arises from trisomy of chromosome 21, but its molecular and pathological consequences are not fully understood. In this study, we compared Dp1Tyb mice, a DS model, against their wild-type (WT) littermates of both sexes to investigate the impact of DS-related genetic abnormalities on the brain phenotype.

We performed *in vivo* whole brain magnetic resonance imaging (MRI) and hippocampal ^1^H magnetic resonance spectroscopy (MRS) on the animals at 3 months of age. Subsequently, *ex vivo* MRI scans and histological analyses were conducted post-mortem. Our findings unveiled distinct neuroanatomical and biochemical alterations in the Dp1Tyb brains.

Dp1Tyb brains exhibited a smaller surface area and a rounder shape compared to WT brains. Regional volumetric analysis revealed significant changes in 26 out of 72 examined brain regions, including the medial prefrontal cortex and dorsal hippocampus. These alterations were consistently observed in both *in vivo* and *ex vivo* imaging data. Additionally, high-resolution *ex vivo* imaging enabled us to investigate cerebellar layers and hippocampal subregions, revealing selective areas of decrease and remodelling in these structures.

An analysis of hippocampal metabolites revealed an elevation in glutamine and the glutamine/glutamate ratio in the Dp1Tyb mice compared to controls, suggesting a possible imbalance in the excitation/inhibition ratio. This was accompanied by the decreased levels of taurine. Histological analysis revealed fewer neurons in the hippocampal CA3 and DG layers, along with an increase in astrocytes and microglia. These findings recapitulate multiple neuroanatomical and biochemical features associated with DS, enriching our understanding of the potential connection between chromosome 21 trisomy and the resultant phenotype.

## 1. INTRODUCTION

Down syndrome (DS) is one of the most common genetic conditions, affecting approximately 1 of 800 new-borns, caused by the presence of an extra copy of human chromosome 21 (Hsa21). DS presents with a wide-ranging and highly variable constellation of characteristics including altered hearing and vision, learning and language problems, congenital heart defects, and an increased risk of developing comorbidities such as diabetes, depression, and Alzheimer’s disease (Antonarakis et al., 2020; Lott, 2012). People with DS can have profound cognitive, executive and memory deficits (Grieco et al., 2015; Jafri and Harman, 2020; Pennington et al., 2003; Tungate and Conners, 2021).

While the underlying changes in the brain are not yet fully understood, advances in imaging and other *in vivo* methods have provided a more detailed picture about structural, functional, and metabolic consequences of DS (Brown et al., 2021; Dierssen, 2012; Jenny A. Klein and Haydar, 2022). To this end, magnetic resonance imaging (MRI) studies have shown decreases in the overall brain volume in people with DS compared to neurotypical people, with several brain regions particularly affected, such as the frontal lobes, the hippocampus, and the cerebellum (Koenig et al., 2021; McCann et al., 2021). On the other hand, the complexity of this syndrome is also reflected in some posterior cortical and subcortical regions (including brainstem) that are relatively preserved or even increased in volume (Pinter et al., 2001). Regarding cellular and molecular changes, increases in glial markers (such as inositol and glutamine) and decreases in neuronal markers (such as glutamate) have been reported using magnetic resonance spectroscopy (MRS), in both humans and animal models of DS (Lin et al., 2016; Patkee et al., 2021; Shonk and Ross, 1995; Vacca et al., 2019). These changes have been explored more in-depth *ex vivo,* with human post-mortem studies confirming an increase in activated astrocytes and microglia in selective brain regions, such as the frontal lobe and the hippocampal dentate gyrus (Chen et al., 2014; Jenny A Klein and Haydar, 2022; Pinto et al., 2020).

Further advancements in our understanding of the “trisomic brain” have allowed us to discern the patterns of abnormalities with greater precision. By investigating various aspects of DS such as age, sex, and environmental factors, we are gaining a more detailed understanding of this condition (Brown et al., 2021; Cañete-Massé et al., 2022; Dierssen, 2012; Koenig et al., 2021). For example, age is a crucial factor influencing the neuroanatomical and neuropsychological observations associated with DS (Dierssen, 2012; Lockrow et al., 2012; Teipel et al., 2004). This is likely associated with the higher increased risk of developing Alzheimer’s disease in DS (Gomez et al., 2020; Hartley et al., 2015; Snyder et al., 2020) – as individuals with DS age, the impact on neuroanatomy and cognitive function becomes more pronounced (Iulita et al., 2022). In addition, biological sex-related factors might also contribute to the observed variations. Notably, females tend to exhibit better cognitive abilities and milder intellectual disability compared to males within the DS population (Aoki et al., 2018; Kittler et al., 2004; Määttä et al., 2006). Sexual dimorphism in DS, also seen in animal models such as the Dp(10)1Yey mice (Block et al., 2015; Hawley et al., 2022; Minter and Gardiner, 2021) echoes other psychiatric and neurodevelopmental disorders including depression, anxiety, attention-deficit and hyperactivity disorder, and autism (Green et al., 2019; Mandy et al., 2012; Rucklidge, 2010). There is certainly a need for a more detailed evaluation of the relationship between genes, sex, and phenotype, which could improve the fit and the precision of potential future therapeutic interventions (de Sola et al., 2015).

The utilization of animal models has been a key in advancing our understanding of the phenotypic characteristics of DS (Muñiz Moreno et al., 2020). However, modelling DS in mice poses challenges due to the dispersion of regions orthologous to Hsa21 across three chromosomes (Mmu10, Mmu16, and Mmu17) in mice (Herault et al., 2017). As a result, accurately modelling the DS condition in mice is complex, and some older models have triplicates of non-DS related gene sequences. Nevertheless, through recent advances in genetic engineering, we have created a more precise model that has an extra copy of most orthologues of Hsa21 genes: theDp1Tyb mouse model carries a duplication from *Lipi* to *Zbtb21* on Mmu16, spanning 23 Mb and 148 coding genes with orthologues on Hsa21 (Lana-Elola et al., 2016). The duplication of these genes leads to multiple DS-like phenotypes, including cardiac defects, learning and memory deficits, and sleep problems (Chang et al., 2020; Lana-Elola et al., 2021, 2016). Importantly, Dp1Tyb mice also have craniofacial dysmorphologies characteristic of DS – such as reduced size of the cranium and mandible, brachycephaly (shortened head), and mid-facial hypoplasia – revealed by recent cranial examination using high-resolution computed tomography imaging (Toussaint et al., 2021). Here, we focused on exploring the macroscopic anatomical, chemical, and cellular changes in Dp1Tyb mouse brains using MR-based techniques and histology, investigating also whether the genotype and sex interact to produce volumetric and cellular brain alterations. Our analysis provides an improved basis for understanding the cognitive impairment and craniofacial abnormalities that we previously observed in this model (Lana-Elola et al., 2021; Toussaint et al., 2021).

## 2. MATERIALS & METHODS

### 2.1. ANIMALS

C57BL/6J.129P2-Dp(16*Lipi*-*Zbtb21*)1TybEmcf mice (hereafter referred to as Dp1Tyb) were generated using long-range Cre/loxP mediated recombination to duplicate the region of Mmu16 from *Lipi* to *Zbtb21*, as previously described (Lana-Elola et al., 2016). All mice were backcrossed to C57BL/6J for at least ten generations, and their genotypes were established using DNA samples isolated from ear biopsies.

The mice were housed in individually ventilated cages of 2-5 age-matched animals, under controlled environmental conditions (24–25°C; 50%–60% humidity; 12 h light/dark cycle) with free access to food and water. We used a total of ten Dp1Tyb mice (5 males and 5 females) and fourteen age-matched wild-type (WT) littermates (8 males and 6 females) of 14 ± 1 weeks of age. Of these, two did not undergo *in vivo* MRI (one male WT and one male Dp1Tyb) due to COVID-19-related restrictions, two Dp1Tyb females were excluded from ex vivo MRI and histology due to a failure of perfusion, and one additional WT male was excluded from histology due to technical problems.

### 2.2. IN VIVO MAGNETIC RESONANCE IMAGING AND SPECTROSCOPY

The *in vivo* scanning was performed in a 9.4 T horizontal bore Bruker BioSpec 94/20 scanner, using an 86-mm volume resonator and a 2×2 phased array surface RF coil. The mice were first anesthetized with a mixture of air with 30% oxygen and 4% isoflurane and then transferred to the scanner bed. The animals’ respiration rate and core temperature were monitored and maintained during scanning at 60-80 breaths/min and 37 ± 0.5°C, respectively, by adjusting the isoflurane level between 2-2.5% and using a circulating hot water system (SA Instruments, Inc).

#### 2.2.1. BRAIN STRUCTURE AND VOLUME

Three-dimensional (3D) T1-weighted images were acquired using an MP2RAGE sequence (Marques et al., 2010), with the following parameters: echo time (TE) = 2.7 ms, repetition time (TR) = 7.9 ms, inversion times (TI) = 800 and 3500 ms, flip angle (FA) = 7°/7°, segment TR = 7000 ms, 1 segment, 4 averages, field of view (FOV) = 17.4×16.2×9.6 mm, matrix = 116×108×64. To combine the complex MP2RAGE images from the four elements in the phased array surface coil, a reference scan was acquired using a 3D ultra-short echo time (UTE3D) sequence: TE = 8.13 μs, TR = 4 ms, FA = 5°, 28733 radial projections, FOV = 35×35× 35 mm, matrix = 96×96×96.

The MP2RAGE complex images were combined using the COMbining Phase data using a Short Echo time Reference scan (COMPOSER) approach (Robinson et al., 2017) implemented in QUantitative Imaging Tools (QUIT, qi composer.sh) (C Wood, 2018). Then, QUIT qi mp2rage was used to create uniform T1-weighted images and T1 relaxation time maps. To assess brain volume and T1 changes, tensor-based morphometry and atlas-based analysis — using a modified version of the Allen Mouse Brain Atlas consisting of 72 regions of interest, (Wang et al., 2020)— were performed on these images as previously described (Bogado Lopes et al., 2023; Mueller et al., 2021).

#### 2.2.2. SINGLE VOXEL ^1^H SPECTROSCOPY

We used single voxel ^1^H spectroscopy (MRS) to quantify alterations in the hippocampal metabolite profiles in Dp1Tyb mice. The individual spectra were acquired using a Point REsolved Spectroscopy (PRESS) pulse sequence (Yahya, 2009) with the following parameters: TE = 8.26 ms, TR = 2500 ms, 512 averages, acquisition bandwidth = 4401 Hz, 2048 acquisition points, voxel size = 2.2 × 1.2 × 2.5 mm. Outer volume suppression and water suppression with variable pulse power and optimized relaxation delays (VAPOR) were used in order to mitigate the contribution of signal from outside the prescribed voxel and suppress unwanted signal from water, as described by (Kiemes et al., 2022).

The MR spectra obtained from each animal were analysed with two software packages: FID Appliance (FID-A) (Simpson et al., 2017) and Linear Combination (LC) Model version 6.3 (Provencher, 2001, 1993). In total, eleven metabolites were quantified, as per (Kiemes et al., 2022): creatine (Cr), gamma-aminobutyric acid (GABA), glutamine (Gln), glutamate (Glu), glutathione (GSH), *myo*-inositol (Ins), lactate (Lac), N-acetyl-aspartate (NAA), phosphocholine (PCh) phosphocreatine (PCr), and taurine (Taur).

### 2.3. EX VIVO MAGNETIC RESONANCE IMAGING

At the end of the *in vivo* MR scan, Dp1Tyb and WT mice were deeply anesthetized with a mixture of drugs (0.05mg/kg Fentanyl, 5mg/kg Midazolam and 0.5mg/kg Medetomidine) and intracardially perfused with phosphate-buffered saline (PBS, pH 7.2), followed by 4% formaldehyde in PBS. The heads of the animals were removed and post-fixed overnight at 4°C in 4% formaldehyde and then immersed in 8 mM Gd-DTPA (Magnevist, Bayer) in PBS + 0.05% sodium azide for at least three months prior to *ex vivo* MRI.

The *ex vivo* scanning was performed on a 9.4T Bruker BioSpec 94/20 with a 39-mm transmit/receive volume coil. The heads were scanned four at a time secured in a custom-made holder and immersed in perfluoropolyether (Galden®, Solvay). A FLASH sequence was used with the following parameters: TE/TR = 6/20 ms, FA = 33°, FOV = 25×25× 20 mm, matrix = 625×625×500, 7 averages, scan time = 14 h.

The images were first processed using the same pipeline as the one employed for the *in vivo* scans, applying tensor-based morphometry and atlas-based regional analysis. In addition, the increased resolution of the *ex vivo* scans (40-μm isotropic) allowed us to perform more detailed analysis to quantify the differences in the structure of the cerebellum and the hippocampus, regions particularly affected in people with DS. We quantified the cerebellar morphometric changes between WT and Dp1Tyb animals in terms of lobular volume and thickness through the analysis framework described by (Ma et al., 2020). This analysis is based on the extraction of the middle Purkinje layer through surface segmentation to estimate the structural morphologies of the granular and molecular layers. Similarly, we analysed the volumes and thicknesses of dorsal and ventral hippocampal regions delineated in the Allen Mouse Brain Atlas: CA1 and CA3 subfields and the molecular, granule cell, and polymorphic layers of the dentate gyrus (DG).

### 2.4. IMMUNOFLUORESCENCE

After *ex vivo* scanning, the brains were extracted, immersed in 30% sucrose for 2 to 3 days, and serially sectioned in 35 µm thick coronal sections using an HM 430 Sliding Microtome (Thermo Fisher Scientific). Two sets of free-floating, double immunofluorescence (IF) staining were performed, each one using 3 to 4 sections per animal (bregma: -1.70 mm to -1.94 mm): the first was to analyse NeuN (neurons) and GFAP (astrocytes), and the second to quantify Iba1 (microglia/macrophages) and SV2A (synaptic density marker).

The sections were heated for 30 min in sodium citrate buffer (pH 6.0) at 80°C, permeabilized for 10 min with 0.3 % Triton ×100 (only for Iba1/SV2A IF), and incubated for 1 h with a blocking solution containing either 10% skim milk powder in tris-buffered saline (TBS) with 0.3% Triton (NeuN/GFAP IF) or 10% donkey serum in TBS with 0.05% Tween x20 (Iba1/SV2A IF). Immediately after, the sections were incubated overnight with the appropriate primary antibodies diluted in blocking buffer, at 4°C (see *Supplementary Table 1*). The next day, the sections were washed and incubated with the secondary antibody for 2 h (NeuN/GFAP IF) or 1.5 h (Iba1/SV2A IF) at room temperature. Finally, the sections were counterstained for 5 min with 300 nM 4′,6-diamidino-2-phenylindole (DAPI), mounted, and coverslipped with antifade medium (FluorSave™, Calbiochem, #345789–20). A negative control (incubated only with the secondary antibody) was used to confirm the primary antibody specificity for each protein.

Three regions of interest (ROIs) were imaged and analysed per section, corresponding to different hippocampal subregions: the pyramidal cell layer of CA1 and CA3, and the polymorphic layer of the DG (see *Supplementary Figure 1*). For each ROI, two to four images were captured at 40× magnification using a Zeiss AxioImager Z1 widefield fluorescence microscope (Carl Zeiss, Ltd), a monochrome AxioCamMR3 camera and the AxioVision 4.8. imaging software (Carl Zeiss, Welwyn, Garden City). This method resulted in representative sampling of the different ROIs, ensuring the reliability of our results.

An average of 32 ± 5 images per animal (2 to 4 images per ROI × 3 ROIs × 3 to 4 slices per animal) were analysed with ImageJ (ver. 1.8.0, NIH, USA). The *Cell Counter* plugin was used to manually quantify the number of cells (astrocytes, neurons, or microglia) per mm^2^, aided by DAPI counterstaining. SV2A-immunoreactivity was quantified by measuring the integrated density after background and non-specific binding subtraction.

### 2.5. STATISTICAL ANALYSIS

Different software packages were employed to statistically analyse differences between Dp1Tyb and WT mice, depending on the modality.

The voxel-wise differences in regional brain volumes (log-transformed Jacobian determinants of the normalisation deformation fields) and T1 relaxation times were analysed using FSL randomise (Winkler et al., 2014). Nonparametric permutation inference was performed using 5000 permutations, threshold-free cluster enhancement, and family-wise error (FWE) correction.

The cerebellar cortical laminar and hippocampal subregional image processing and groupwise surface-based morphological statistical analysis were achieved through the Multi Atlas Segmentation and Morphometric Analysis Toolkit (MASMAT) (Ma et al., 2014)^1^ and the Shape & Morphological Analysis and Rendering Toolkit (SMART) (Ma et al., 2020) ^2^ accordingly.

SPSS (IBM®SPSS® Statistics 26; USA) was used to analyse the atlas-based regional changes in volume and T1 relaxation time, hippocampal metabolites, and IF. Firstly, the Shapiro–Wilk test was used to test the data for normality and the Levene’s test was employed to assess the homogeneity of variances. Then, a two-way (genotype × sex) ANOVA was performed for all data, setting the threshold of statistical significance at p = 0.05. Finally, we corrected for multiple comparisons with the “two-stage” Benjamini, Krieger and Yekutieli procedure (*q-value* set as 0.05) with GraphPad Prism (version 8), controlling the false discovery rate (FDR) (Benjamini et al., 2006). Additionally, a potential correlation between the hippocampal volume, the expression of different metabolites (MRS), and the cellular and synaptic density in different hippocampal subregions (IF) was explored with SPSS though the one-tailed Pearson’s *r* correlation coefficient.

Finally, GraphPad Prism (version 9.4.1) was used to graphically represent the results, expressed as mean ± SEM.

## 3. RESULTS

### 3.1. IN VIVO MAGNETIC RESONANCE IMAGING AND SPECTROSCOPY

#### 3.1.1. BRAIN STRUCTURE AND VOLUME

##### 3.1.1.1. Whole brain structure and volume

The brains of people with DS exhibit differences in shape and structure, which led us to assess the global characteristics of the Dp1Tyb brains, comparing male and female mutant mice with WT animals (*Figure 1A*).

**Figure 1.**
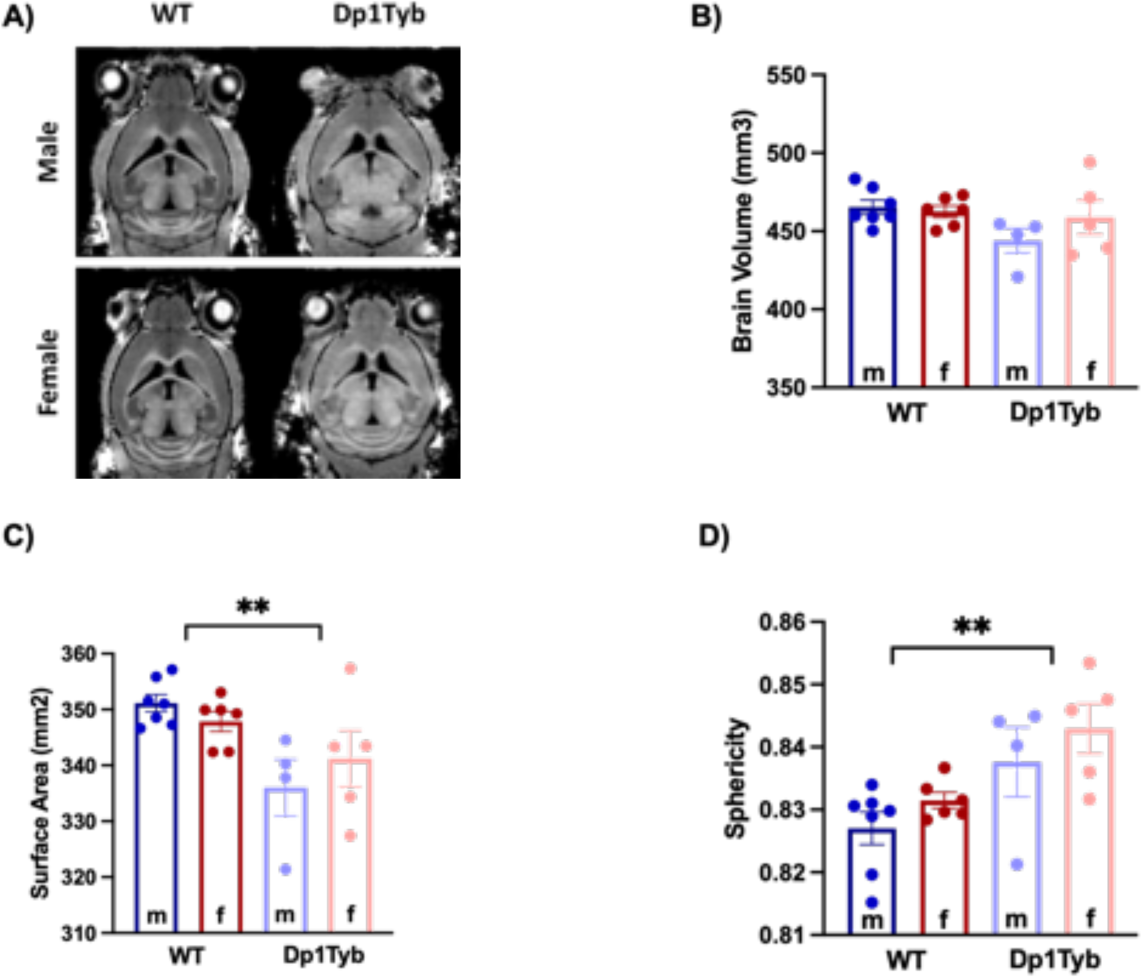
Brain characteristics of wild-type (WT) and Dp1Tyb mice. A) Representative structural T1-weighted brain images. B - D) Quantification of brain volume, surface area, and sphericity of WT (males = 7, females = 6) vs Dp1Tyb (males = 4, females = 5). Bars represent the mean ± SEM. (**) indicates significant differences between WT and Dp1Tyb mice in brain surface area and sphericity (p < 0.01) as yielded by a two-way ANOVA (genotype × sex). m: male, f: female.

Overall, the brains of Dp1Tyb mice had a smaller surface area (F_1,18_ = 11.12; p = 0.004) and were significantly rounder (F_1,18_ = 11.31; p = 0.003) than the brains of WT animals (*Figure 1A, C, D)*. These differences can be seen when looking at the representative extracted brains, as shown in *Supplementary Figure 2*. Whole brain volumes measured *in vivo* were not significantly different between the genotypes, however there was a significant genotype effect when the volume of the brains was quantified *ex vivo* (*Supplementary Table 2*).

No significant differences due to sex or sex × genotype interaction were found (n.s., p > 0.05).

##### 3.1.1.2. Differences in regional brain volumes

We next performed a voxel-based analysis to detect the areas of the brain that are driving the observed global differences between Dp1Tyb and WT littermates. Dp1Tyb mice showed significant decreases in the volume of multiple regions (blue colours, Figure 2). However, other regions were significantly smaller in Dp1Tyb (red colours, Figure 2), in particular the brainstem.

**Figure 2.**
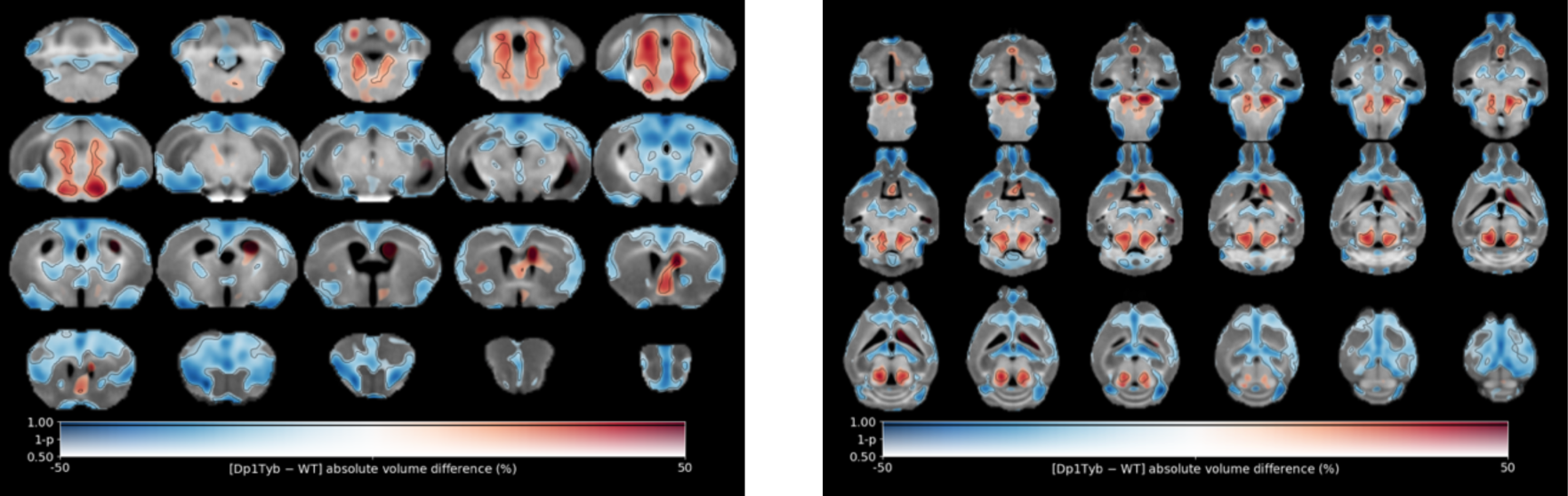
Voxel-wise differences in volume between Dp1Tyb and WT mice. Map of voxel-wise differences in volume between Dp1Tyb (n = 9) and WT mice (n = 13), derived from in vivo MR images and overlaid on the T1-weighted study-specific template. The map is displayed in the coronal plane (left image, caudal-rostral) and the horizontal plane (right image, ventral-dorsal). The colour of the overlay indicates the percent volume difference (cool colours indicate reduced volume and hot colours increased volume in Dp1Tyb compared to WT mice), and the opacity of the overlay indicates the significance of the volume difference (regions where the FWE-corrected p > 0.5 are completely transparent, and regions where the FWE-corrected p = 0 completely opaque). Clusters where the FWE-corrected p < 0.05 are contoured in black.

The analysis of sex-related differences (all males vs all females) revealed several cortical clusters with an increased volume in the brain of females (*see Supplementary Figure 3*). However, no significant genotype × sex interaction was found.

Subsequently, we explored how the changes observed in individual voxels correspond to volume alterations in specific brain regions. To that end, we quantified the volumes of 72 regions of interest (*ROIs*) derived from the Allen mouse brain atlas. The statistical analysis highlighted significant differences between WT and Dp1Tyb mice in 26 out of 72 regions (*Figure 3A*). These regions could be clustered according to their role in biological processes (*Supplementary Table 3*). For example, Dp1Tyb mice have significant decreases in the volumes of regions involved in decision-making and executive processes (e.g. orbital and medial prefrontal cortices) and in working memory and spatial memory tasks (e.g. retrosplenial and medial prefrontal cortices, and dorsal hippocampus). Other regions with significantly decreased volume included those related to processing of sensorial stimuli (e.g. auditory and olfactory cortex) and emotions (e.g. insular and cingulate cortices and amygdala), as well as regions implicated in generating a behavioural response to stress and anxiety (e.g. habenula and dorsal peduncular area). However, we observed a significant increase in the volume of regions involved in the regulation of the sleep-wake cycle and various autonomic functions (such as the pons and different brainstem nuclei). These results are presented in *Figure 3B*.

**Figure 3.**
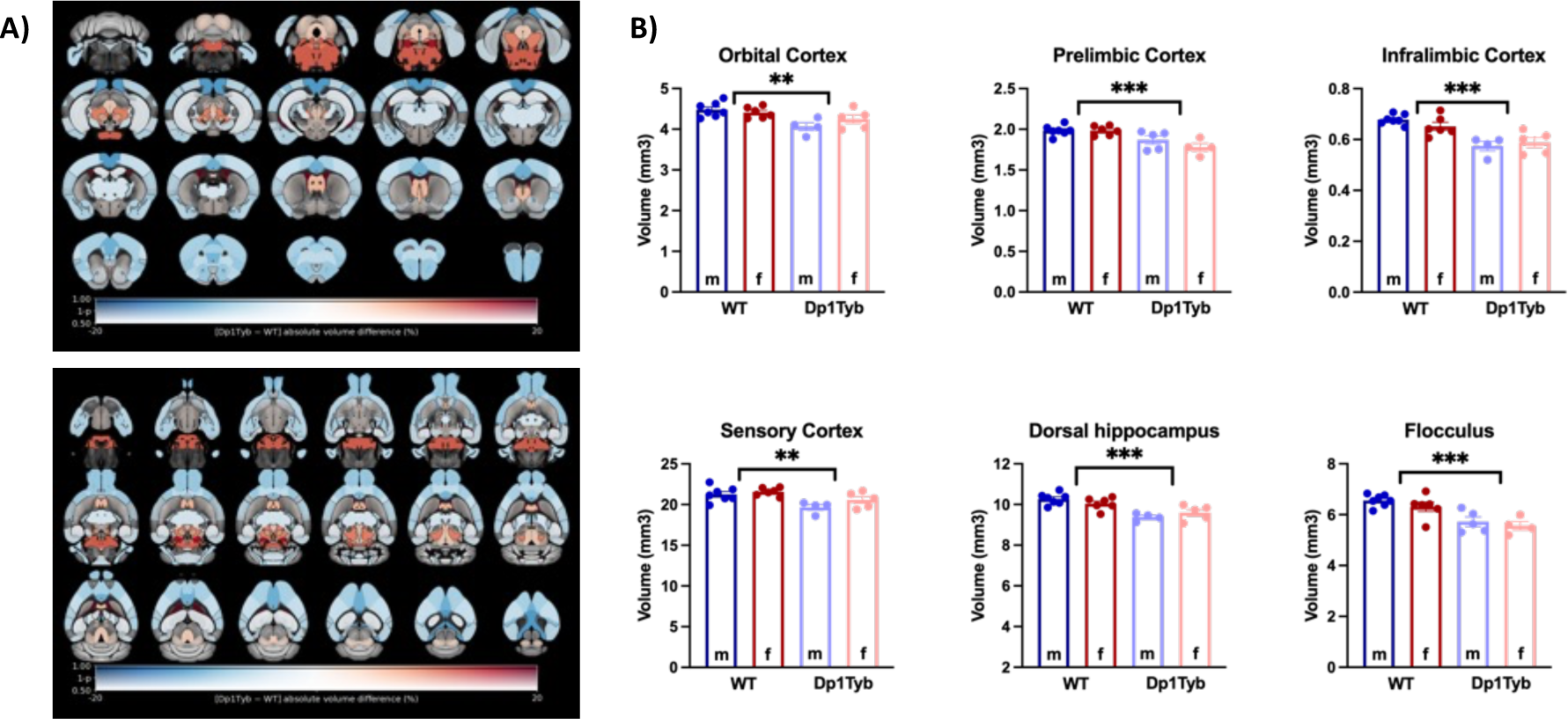
Differences in regional volumes between Dp1Tyb and WT mice. **A)** Maps of differences in regional volumes between Dp1Tyb (n = 9) and WT mice (n = 13), calculated from in vivo MR images and overlaid on the Allen mouse brain template (72 ROIs). The map is displayed in the coronal plane (top image, caudal-rostral) and the horizontal plane (bottom image, ventral-dorsal). The colour of the overlay indicates the percent volume difference (cool colours indicate reduced volume in Dp1Tyb compared to WT mice), and the opacity of the overlay indicates the significance of the volume difference (regions where the FDR-corrected p > 0.5 are completely transparent, and regions where the FDR-corrected p = 0 completely opaque). ROIs for which the FDR-corrected p < 0.05 are contoured in black. **B)** Selection of ROIs with significant differences in volume between WT and Dp1Tyb brains. The plots display the mean ± SEM for each group. Group comparisons were performed with a two-way ANOVA (genotype × sex), using the FDR to correct for multiple comparisons (Q = 5%). The effect of genotype is represented as: **p < 0.01, ***p < 0.001.

Significant sex differences (females presenting bigger volume than males) were only found in two regions — the auditory cortex and the claustrum — and a significant genotype × sex interaction in only one region — the postrhinal cortex (no significant difference in volume in WT males vs Dp1Tyb males, but bigger volume in Dp1Tyb females compared to WT females - F_1,18_ = 9.70; p = 0.006).

#### 3.1.2. CHANGES IN T1 RELAXATION TIME

T1 relaxation time provides information about tissue water content and lipid concentration (indirect measure of axonal organisation and myelin production), and it is considered an optimal marker of brain maturation (Nossin-Manor et al., 2013; Schneider et al., 2016).

The analysis of T1 relaxation maps derived from the MP2RAGE images revealed a subtle yet global reduction of T1 relaxation time in the Dp1Tyb brains. As shown in *Figure 4A*, large areas were affected, some of which were colocalised in areas that also presented a volume decrease, such as the prelimbic cortex and the flocculus (see *Figure 4B*). However, neither voxel-wise nor atlas-based regional analysis showed significant differences in T1 after correcting for multiple comparisons (no significant effect of sex, genotype, or sex × genotype).

**Figure 4.**
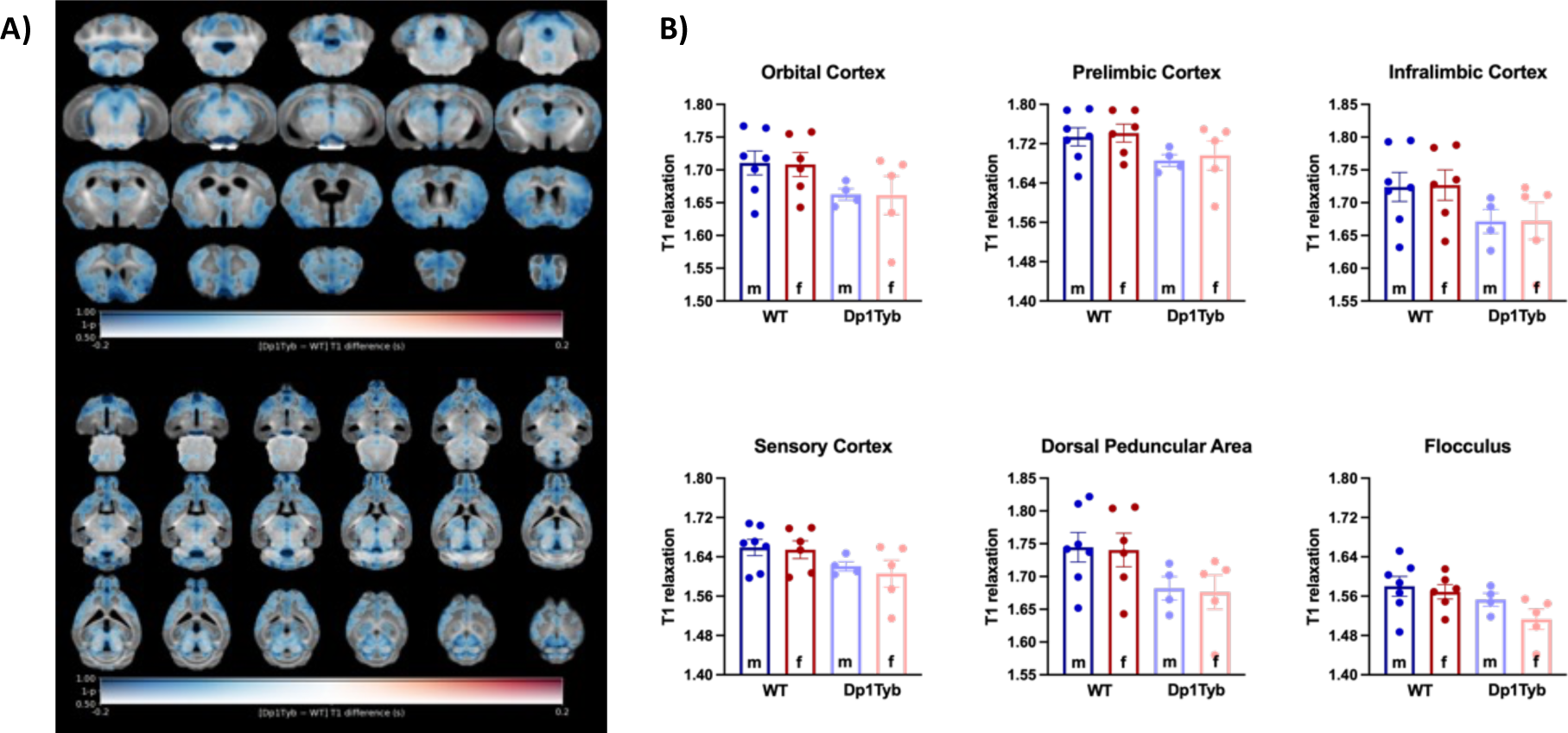
Differences in T1 relaxation time between Dp1Tyb and WT mice. **A)** Maps of voxel-wise differences between Dp1Tyb (n = 9) and WT mice (n = 13), calculated from in vivo MR images and overlaid on the T1-weighted study-specific template. The map is displayed in the coronal plane (top image, caudal-rostral) and the horizontal plane (bottom image, ventral-dorsal). The colour of the overlay indicates the difference in T1 (cool colours indicate reduced T1 in Dp1Tyb compared to WT mice), and the opacity of the overlay indicates the significance of the difference (regions where the FWE-corrected p > 0.5 are completely transparent, and regions where the FWE-corrected p = 0 completely opaque). None of the clusters were significantly different at the level of FWE-corrected p < 0.05. **B)** T1 relaxation times of example ROIs. The plots display the mean ± SEM for each group. The effects of genotype, sex, and the interaction between both variables were not significant according to a two-way ANOVA (genotype × sex), with p < 0.05.

#### 3.1.3. SINGLE VOXEL ^1^H SPECTROSCOPY

We next assessed biochemical alterations in Dp1Tyb mice, comparing the concentrations of hippocampal metabolites with WT animals using a two-way (genotype × sex) ANOVA. Absolute values of all metabolites, as well as glutamine/glutamate ratio are shown in Supplementary Table 4. While there were trend differences in the concentration of several metabolites (including glutamate, glutathione, lactate and N-acetyl-aspartate) only three metabolites remained significant when data were corrected for multiple comparisons based on 11 metabolites. These were significant increase in the concentration of glutamine (Gln: F_1,18_ = 15.47; p = 0.001) and the glutamine/glutamate ratio (Gln/Glu: F_1,18_ = 14.22; p = 0.001), and a significant decrease in the concentration of taurine (Tau: F_1,18_ = 22.51; p < 0.001) in Dp1Tyb compared to WT animals *(Figure 5)*. Metabolite concentrations were not significantly different between males and females, and there was no statistically significant genotype × sex interaction.

**Figure 5.**
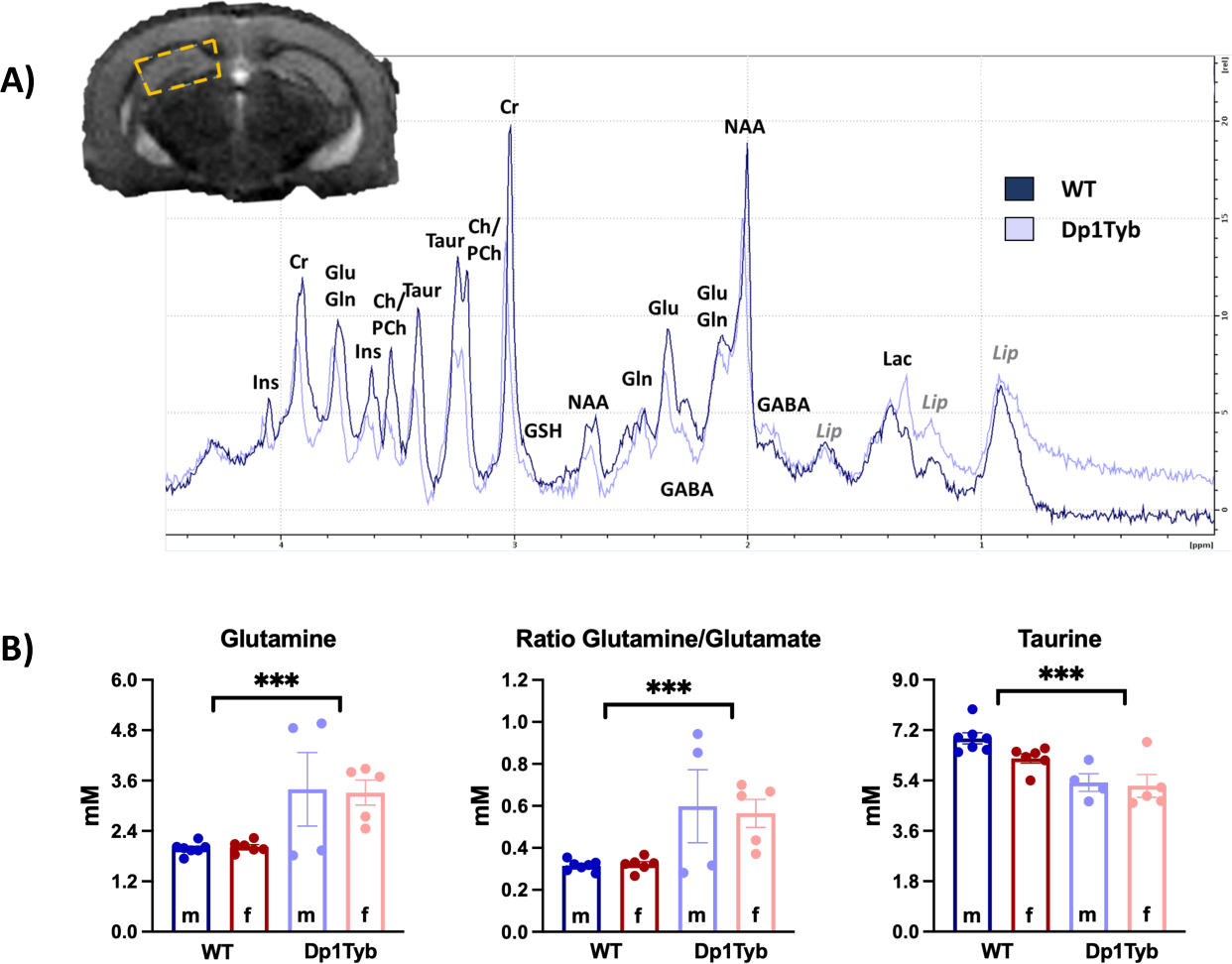
Differences in the MR spectra between Dp1Tyb (n=9) and WT (n=13) mice. A) Location of the MRS voxel in the hippocampus (yellow box) and representative spectra from one WT and one Dp1Tyb mouse. B) The plots represent the mean ± SEM. Group differences were determined by a two-way ANOVA (genotype × sex), using the FDR to correct for multiple comparisons (Q = 5%). The effect of genotype is represented as ***p < 0.001. Abbreviations: Cr (creatine), GABA (gamma-aminobutyric acid), Gln (glutamine), Glu (glutamate), GSH (glutathione), Ins (myo-inositol), Lac (lactate), Lip (Lipids), NAA (N-acetyl-aspartate), PCh (phosphocholine), Taur (taurine).

### 3.2. EX VIVO MAGNETIC RESONANCE IMAGING

Following *in vivo* scanning, the mice underwent perfusion, and the fixed heads were doped with a gadolinium-based contrast agent before being imaged *ex vivo* at higher resolution. Employing a similar analysis pipeline used for the *in vivo* scans, we observed a consistent pattern of voxel-wise differences between Dp1Tyb and WT mice. Several regions in the Dp1Tyb brains exhibited decreased volume, including within the orbital, prelimbic, motor, and piriform cortices. Conversely, other regions displayed increased volume, such as the septal nucleus, diagonal band, and various pontine nuclei (*Supplementary Figure 4*).

The observed changes were more distinct and finely detailed in the *ex vivo* scans compared to the *in vivo* scans. This likely stems from the higher resolution of the *ex vivo* scans, as well as the physical alterations caused by factors such as death and perfusion (Holmes et al., 2017). Moreover, *ex vivo* images revealed specific differences in the layers of structures with distinct layers, such as the hippocampus and cerebellum. Considering the well-known involvement of the cerebellum and hippocampus in DS, we further investigated these changes by analysing cerebellar and hippocampal substructures using our recently developed analysis pipeline (Ma et al., 2020).

Although the volume of the whole cerebellum was not significantly different between the genotypes *in vivo*, Dp1Tyb mice showed smaller absolute (but not relative) cerebellar volumes when measured *ex vivo* (*F_1,18_ = 6.058; p = 0.0242)* (*Supplementary Table 2*). This appeared to be driven by a decrease in the volume of the granular (*F_1,18_ = 11.74; p = 0.003*) but not molecular layer (*Figure 6A*). There was no significant group difference in the thickness of either layer. Further regional analyses pointed out specific volume decreases in the granular layer lobules 3, 8, 9 and 10, the paraflocculus, and the flocculus (*Figure 6C*). In addition, there were specific decreases in the molecular layer lobules 3 and 8 and in the flocculus (*Supplementary Table 5*).

**Figure 6.**
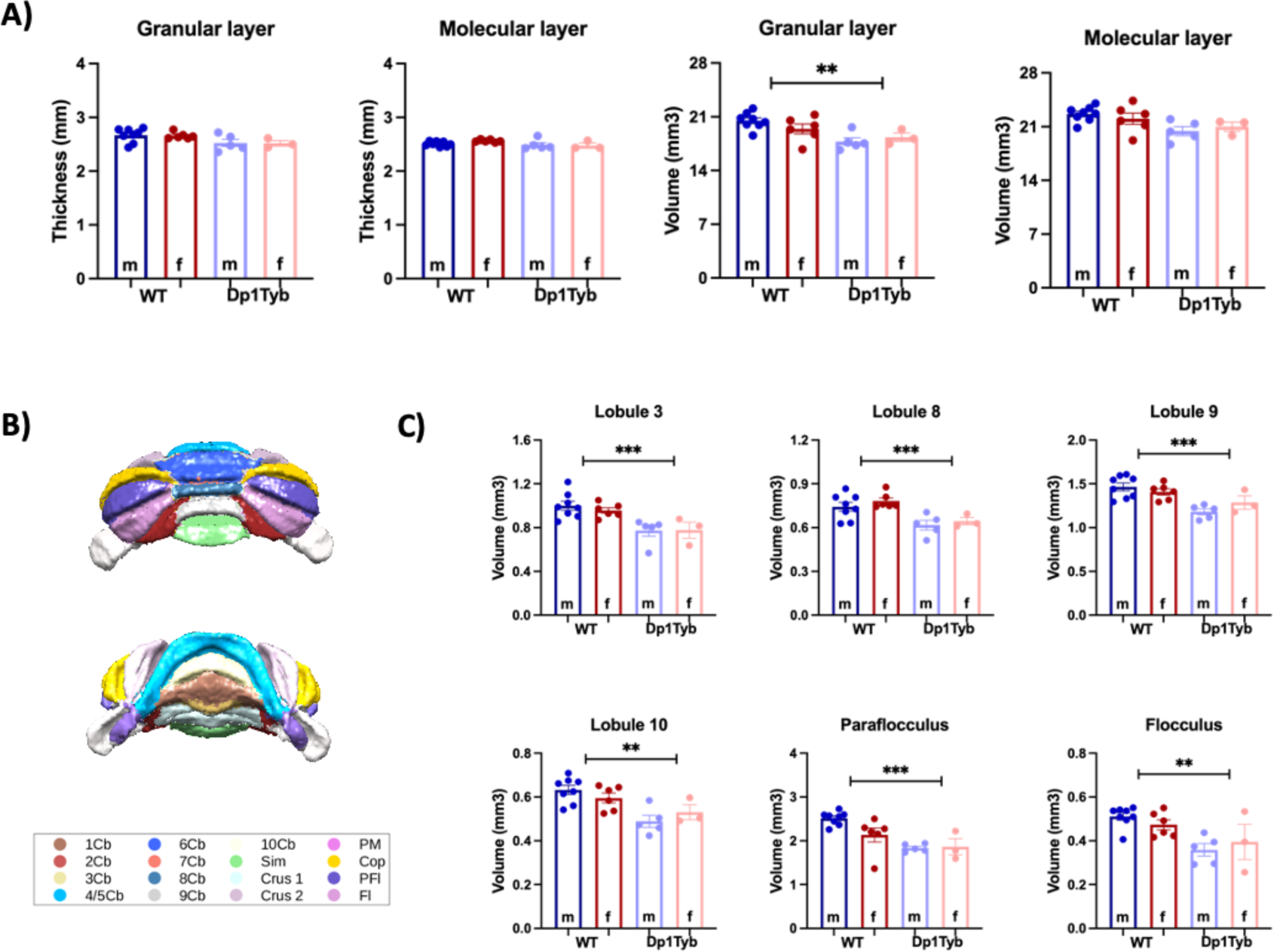
Ex-vivo differences in cerebellar volume between Dp1Tyb and WT mice. A) Quantification of grey matter thickness and volume in Dp1Tyb mice compared to WT. B) Division of cerebellar lobules. C) Cerebellar lobes with significant group differences in the volume of the granular layer. The plots display the mean ± SEM for each group (WT = 14, Dp1Tyb = 8). Group comparisons were performed with a two-way ANOVA (genotype × sex) using the FDR (Q = 5%) to correct for multiple comparisons. The effect of genotype is represented as: **p < 0.01, ***p < 0.001 Abbreviations: lobules of the cerebellar vermis: 1Cb (lobule 1), 2Cb (lobule 2), 3Cb (lobule 3), 4/5Cb (lobule 4/5), 6Cb (lobule 6), 7Cb (lobule 7), 8Cb (lobule 8), 9Cb (lobule 9), 10Cb (lobule 10); lobules of cerebellar hemispheres: Sim (simple lobule), Crus 1 (Crus 1 of the ansiform lobule), Crus 2 (Crus 2 of the ansiform lobule), PM (paramedian lobule), Cop (Copula of the pyramis), PFI (Paraflocculus), FI (Flocculus).

No significant effect of sex or interaction between genotype × sex was found in any of the statistical analyses performed.

The analyses of the hippocampal structures highlighted significant changes in dorsal, but not ventral subregions, in agreement with the results observed *in vivo*. For example, Dp1Tyb mice showed an increase in the thickness of CA3 (F_1,18_ = 7.49; p = 0.014) and a decrease in the volume of CA1 (F_1,18_ = 5.74; p = 0.027) and the molecular layer of DG (MoDG, F_1,18_ = 20.50; p < 0.001) compared to WT mice (see *Figure 7*).

**Figure 7.**
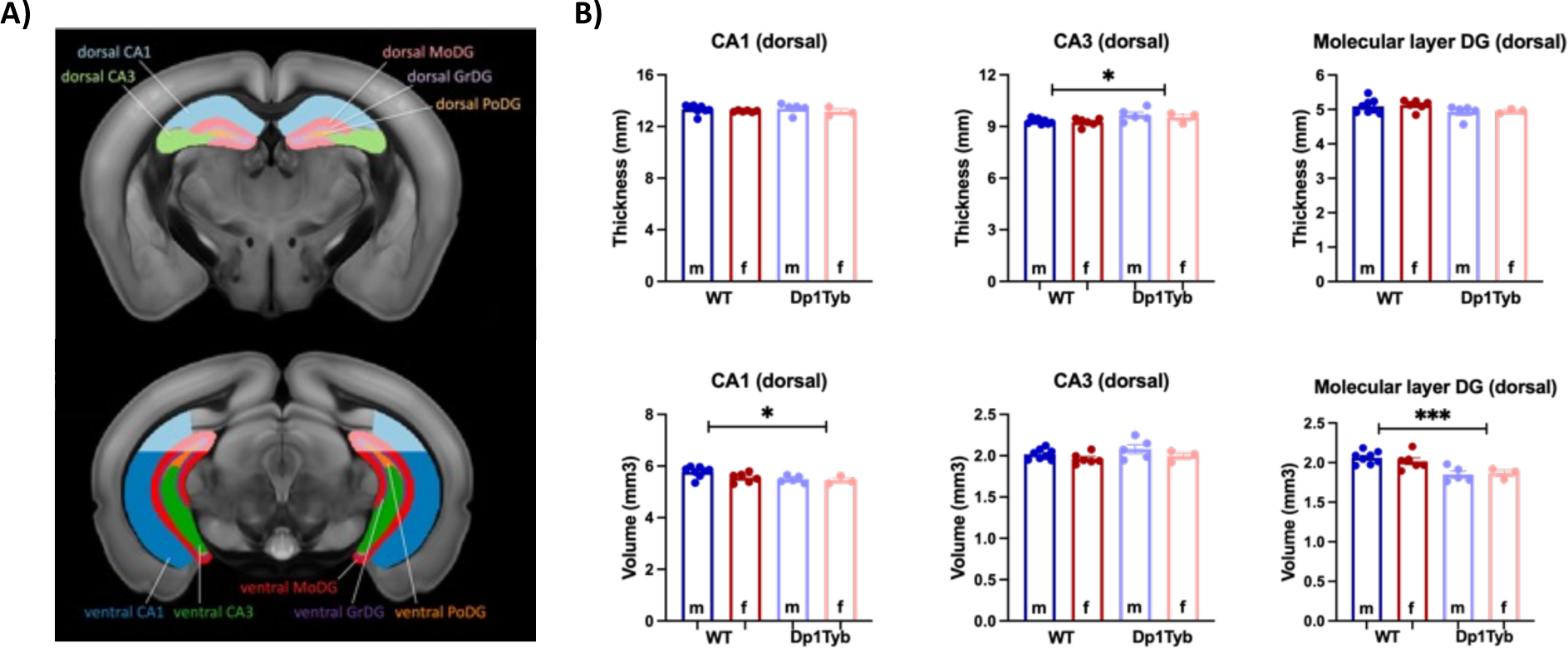
Ex-vivo differences in hippocampal thickness and volume between Dp1Tyb and WT mice. A) Division of dorsal and ventral hippocampal subregions. B) Plots represent hippocampal subregions with significant differences in thickness (dorsal CA3) or volume (dorsal CA1 and MoDG) between WT and Dp1Tyb brains. The plots display the mean ± SEM for each group (WT = 14, Dp1Tyb = 8). Group comparisons were performed with a two-way ANOVA (genotype × sex) using the FDR (Q = 5%) to correct for multiple comparisons. The effect of genotype is represented as: *p < 0.05, ***p < 0.001.

### 3.3. IMMUNOFLUORESCENCE

At the end of the *ex vivo* scans all the brains were extracted, and free-floating IF-based measurements of relevant markers of cells and synapses was conducted.

#### 3.3.1. Number of cells: neurons, microglia and astrocytes

The quantification of the number of hippocampal cells was performed by immunofluorescence staining using markers of neurons (NeuN), astrocytes (GFAP) and microglia (Iba1) (*Figure 8A*).

**Figure 8.**
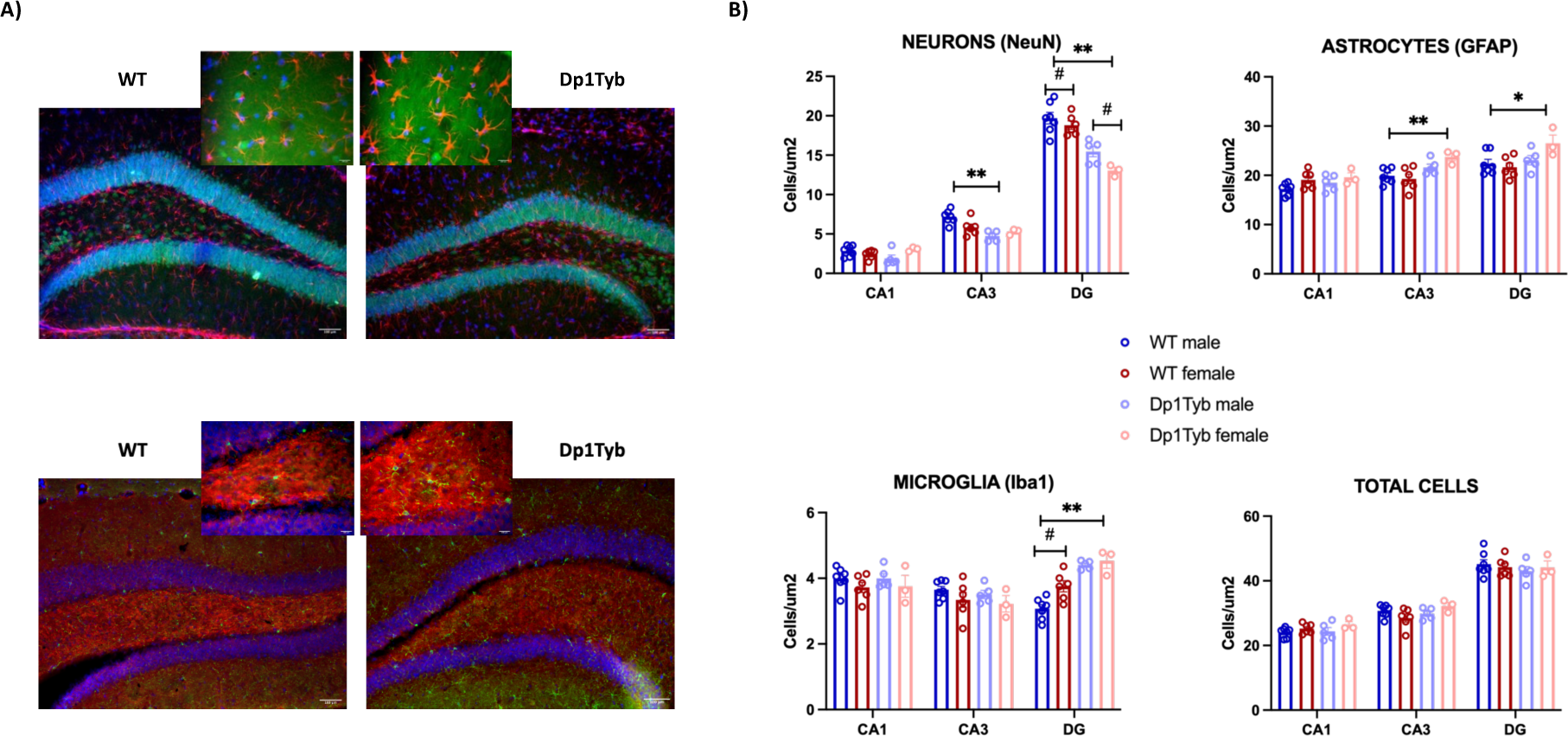
Cell density in hippocampal subregions of Dp1Tyb and WT mice. **A)** Representative widefield fluorescence hippocampal images of WT and Dp1Tyb animals (20x and 40x). The upper images show GFAP (red), NeuN (green), and DAPI (blue), and the bottom images show SV2A (red), Iba1 (green), and DAPI (blue). **B)** Bar plots represent the mean ± SEM for WT (n = 13) and Dp1Tyb mice (n = 7). Statistically significant group differences were assessed by two-way ANOVA (genotype × sex, p < 0.05), with (*p < 0.05 and **p < 0.01) for genotype differences and (# p < 0.05) for sex differences. Abbreviations: CA = cornu ammonis; DG = Dentate gyrus.

Neuronal staining with NeuN showed that Dp1Tyb mice have significantly fewer neurons than WT animals in CA3 (F_1, 17_ = 16.36; p = 0.001) and DG (F_1, 17_ = 49.28; p = 0.0001). In DG, there were also differences between males and females (F_1, 17_ = 5.35; p = 0.035), with females having fewer neurons. Additionally, we observed a significant genotype × sex interaction in CA1 (F_1, 17_ = 6.59; p = 0.020) and CA3 (F_1, 17_ = 6.48; p = 0.021), with Dp1Tyb males having significantly fewer neurons than WT males while there was no significant difference in neuronal numbers between females Dp1Tyb and WT.

Regarding the number of glial cells, GFAP staining showed that Dp1Tyb mice have more astrocytes in both CA3 (F_1, 17_ = 14.23; p = 0.002) and DG (F_1, 17_ = 7.15; p = 0.016), compared with WT animals. In addition, Dp1Tyb mice have more microglial cells (Iba1 staining) in DG (F_1, 17_ = 47.77; p < 0.0001) than WT animals. In this region, there was also a significant effect of sex (females express more GFAP in DG than males — F_1, 17_ = 7.01; p = 0.017), but there was no significant genotype × sex interaction.

Interestingly, the analysis of the “total number of cells” (considered as the sum of the three types of cells) did not to show any differences between the genotypes or sexes, nor was there an interaction between these two variables (*Figure 8B, total cells*).

We next correlated these cell densities with the concentration of metabolites and the volume of the dorsal hippocampus, measured *in vivo* by MRS and MRI, respectively. The average number of hippocampal neurons was positively correlated with the concentrations of taurine (r = 0.796, p < 0.0001) and glutamate (r = 0.414, p = 0.035) and negatively correlated with lactate (r = - 0.554, p = 0.006). Furthermore, the number of hippocampal neurons was also positively correlated with the volume of the dorsal hippocampus (r = 0.704, p < 0.001). The average number of hippocampal astrocytes was positively correlated to glutamine (r = 0.432, p = 0.029) and the glutamine/glutamate ratio (r = 0.396, p = 0.042), and negatively correlated with the volume of the dorsal hippocampus (r = -0.417, p = 0.038). There was no significant correlation between microglia and any MRS/MRI measure.

#### 3.3.2. Hippocampal synaptic density: SV2A

There were no overall differences in SV2A expression between the genotypes in any measured area. However, we did detect a prominent effect of sex (*Figure 9*). Hippocampal SV2A signal was overall lower in females than in males in CA1 (F_1, 17_ = 28.55; p < 0.001), CA3 (F_1, 17_ = 13.01; p = 0.002), and DG (F_1, 17_ = 17.21; p = 0.001) (*Figure 9B*).

**Figure 9.**
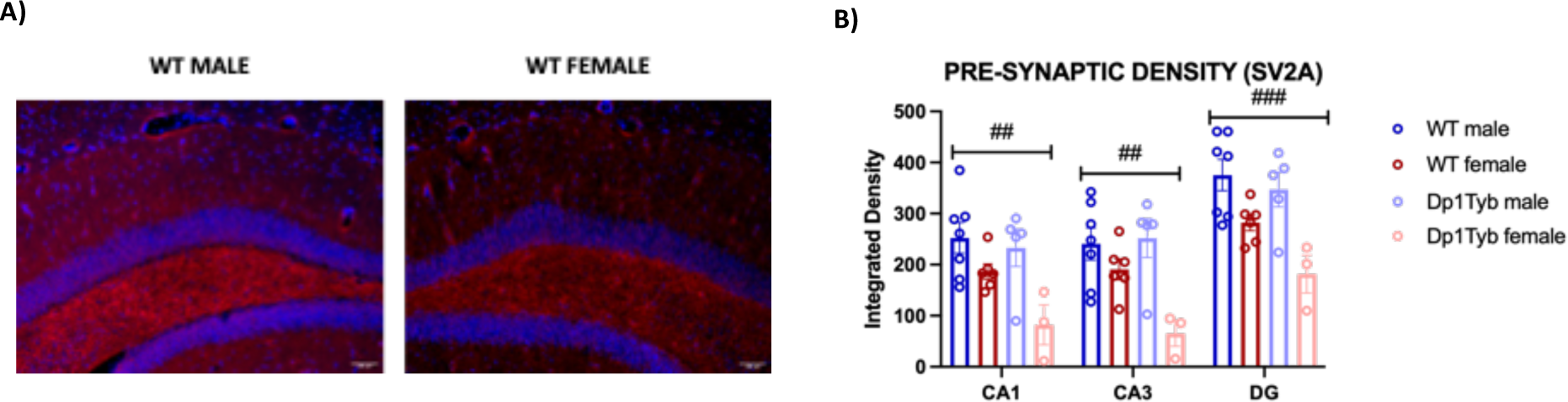
Synaptic density quantified as SV2A expression in hippocampal subregions of Dp1Tyb and WT mice. **A)** Representative widefield fluorescence hippocampal images of a WT male (left) and a WT female (right) (20x). **B)** Bar plots represent the mean ± SEM for WT (n = 13) and Dp1Tyb mice (n = 7). Group differences were determined by a two-way ANOVA (genotype × sex, p < 0.05), with (#) indicating sex-based differences. ## p < 0.01, ### p < 0.001. Abbreviations: CA = cornu ammonis; DG = Dentate gyrus.

Further analyses did not show any significant correlation between hippocampal SV2A expression and any of the eleven hippocampal metabolites quantified (0.061 < p < 0.357), nor between SV2A and the volume of the dorsal hippocampus (r = 0.315, p = 0.095).

## 4. DISCUSSION

This study presents a comprehensive *in vivo* (MRI and MRS) and *ex vivo* (MRI and histology) characterization of brain changes observed in the Dp1Tyb mouse model of DS. We demonstrate that, although the *in vivo* total brain volume of Dp1Tyb mice was not significantly different, the Dp1Tyb brains exhibited a rounder shape and a significantly reduced surface area, which closely resembles findings observed in humans with this condition (McCann et al., 2021). More detailed regional analysis showed significant changes in 26 of the 72 examined regions, most of which were smaller in Dp1Tyb mice but with some notable increases in the subcortical areas of the brainstem. These results are consistent across *in vivo* and *ex vivo* images. The latter allowed us to examine changes in cerebellar layers and hippocampal subregions, two structures known to be particularly affected in DS (Aylward et al., 1999; Pinter et al., 2001; Raz et al., 1995). We additionally sought to examine brain-wide T1 relaxation time, for its known ability to inform about brain maturation, myelination, and integrity of the brain tissue (Tang et al., 2018). We did not see any significant differences in T1 relaxation time in Dp1Tyb mice, except for a non-significant yet widespread trend. Additionally, the quantification of hippocampal metabolites revealed increases in glutamine and glutamine/glutamate ratio and decreases in taurine. Subsequent *ex vivo* histological analysis demonstrated a reduced number of CA3 and DG hippocampal neurons, which seems to be accompanied by an increase in astrocytes and microglia, as the total number of cells did not differ.

Previous imaging studies using computed tomography have identified craniofacial dysmorphologies — such as brachycephaly —in the Dp1Tyb mice and other DS models including the Dp(16)1Yey mice (Lana-Elola et al., 2021; Starbuck et al., 2014; Toussaint et al., 2021). These results, which involve measurements of bone structure and density, faithfully recapitulate aspects of craniofacial alterations observed in humans with DS (Allanson et al., 1993; Korenberg et al., 1994; Richtsmeier et al., 2000). Our study confirms and builds upon these previous findings. Notably, we observed shape differences in the brains of Dp1Tyb mice, presumably reflecting abnormalities of the cranium. The utilization of structural MRI enabled us to conduct a more comprehensive analysis of the internal structure of Dp1Tyb brains. In this regard, 26 anatomical regions showed a change in volume compared to WT mice – reductions for the most part, although some increases were observed too.

Several neuroanatomical changes observed in the Dp1Tyb mouse model may provide insights into the underlying mechanisms of the characteristic phenotype displayed by this model (Chang et al., 2020; Lana-Elola et al., 2021). Notably, we observed a decreased volume of the prefrontal lobe and hippocampus, along with an increased volume of the brainstem. These findings might have implications for understanding the cognitive and behavioural alterations observed in this model. For instance, the decreased volume in regions such as the orbital, prelimbic, and infralimbic cortices might correspond to the reported slower decision-making abilities in Dp1Tyb mice (Chang et al., 2020). Furthermore, the smaller volume of the retrosplenial cortex and dorsal hippocampus aligns with the reported memory deficits (Chang et al., 2020), while the increased volume of brainstem regions involved in the regulation of the sleep-wake cycle might have relation to the disrupted sleep patterns reported in Dp1Tyb mice (Lana-Elola et al., 2021). These observed changes in the Dp1Tyb model closely resemble the heterogeneous findings from humans with DS. For instance, studies in humans have also highlighted the significance of decreased prefrontal cortex and hippocampal volume as key neuroanatomical correlates of cognitive deficits in this population (Fukami-Gartner et al., 2023; McCann et al., 2021; Pinter et al., 2001; Teipel et al., 2004). On the other hand, reports of slight enlargements of deep grey matter structures are reminiscent of our findings of increased volumes of brainstem and septum (Raz et al., 1995; White et al., 2003; Wilson et al., 2019). The biological interpretation of these enlargements, whether they are developmental or compensatory, is not clear due to the diverse range of functions associated with these regions. Overall, our mouse findings further underscore the relevance and validity of the Dp1Tyb model as it recapitulates neuroanatomical alterations observed in humans with DS (Jenny A. Klein and Haydar, 2022).

Previous studies have reported the presence of cerebellar hypoplasia in individuals with DS (Pinter et al., 2001; Weis et al., 1991; Winter et al., 2000) as well as in certain animal models (Kazuki et al., 2022). However, in some cases, these changes were only observed when normalising the cerebellar volume to the total brain or intracranial volume, indicating *relative* volume differences (Ma et al., 2020; Powell et al., 2016). To address this matter, our study comprehensively measured both absolute and relative volumes of the cerebellum, using *in vivo* and *ex vivo* imaging approaches. In line with the findings in human with DS, we observed decreased absolute cerebellar volumes in Dp1Tyb mice, with statistical significance achieved in the *ex vivo images*. This could be attributed to the higher resolution of the *ex vivo* images, facilitating better image registration. Given the intricate morphology of the cerebellum and the varying involvement of its different layers in diverse cognitive functions (Buckner, 2013; Sudarov and Joyner, 2007), we conducted additional in-depth analysis by leveraging the high resolution and enhanced contrast obtained through gadolinium-enhanced *ex vivo* imaging of the brains (Ma et al., 2020). Through these analysis, we observed further *regional* cerebellar volume reductions in the Dp1Tyb mice that were concentrated in the granular layer. This layer, composed of excitatory granule cells and inhibitory Golgi and Lugaro interneurons (Roostaei et al., 2014), is thought to be involved in motor coordination as well as in some non-motor behaviours such as reward expectation-related activity (D’Angelo, 2013; Lackey et al., 2018). The decrease in the volume of this layer could be related to impaired motor function and coordination observed in this model, which was not attributed to altered muscle tone (Lana-Elola et al., 2021). These findings are also consistent with previous MRI and histological analysis performed in other mouse models such as the Tc1 (Ma et al., 2020), as well as in human foetuses with DS (Guidi et al., 2011).

In addition to investigating volumetric changes, we explored potential alterations in T1 relaxation time as an indicator of brain composition. Considering the established utility of T1 relaxation time in assessing brain maturation and neurodegenerative conditions such as Parkinson’s and Alzheimer’s disease (Eriksson et al., 2007; Tang et al., 2018), we anticipated observing changes in T1 relaxation time in Dp1Tyb mice. However, our study revealed only a non-significant, yet widespread decrease in T1 relaxation time. Interestingly, this decrease was consistently distributed bilaterally and aligned with the plausible anatomical regions. While this subtle decrease is not statistically significant, it warrants further study with larger groups of animals, and/or in older age or in models combining DS with Alzheimer’s pathology where the confluence of T1 and neurodegeneration might be more pronounced (Farrell et al., 2022).

From a neurochemical standpoint, we detected significant changes in the concentration of multiple hippocampal metabolites within the Dp1Tyb mice. Amongst the most robustly affected was glutamine. The increased glutamine resulted in a significant increase in the glutamine/glutamate ratio, which could potentially indicate an imbalance in excitatory and inhibitory signalling (E/I balance) – an aspect that has been suggested to be involved in DS (Hamburg et al., 2019). Further research should be focused on unravelling the underlying mechanisms responsible for these potential changes in the E/I balance in DS, as they could serve as useful biomarkers of therapeutic interventions in this mouse model.

We also observed that Dp1Tyb mice have a significant decrease in taurine, a neurotrophic factor involved in brain development. This finding aligns with limited evidence suggesting a similar decrease in taurine in human with DS (Whittle et al., 2007) and there is even anecdotal evidence supporting the potential usefulness of taurine supplementations in DS (Rafiee et al., 2022; Singh et al., 2023). Notably, taurine is found to be decreased during ageing (Singh et al., 2023), and taurine depletion was observed in plasma from AD patients (Rafiee et al., 2022). Collectively, these findings imply a significant involvement of taurine in the neurodevelopmental alterations associated with DS. Additionally, taurine is known to play a role in osmoregulation and may be involved in mitochondrial dysfunction and neuroinflammation (Rafiee et al., 2022), both processes thought to be implicated in DS (Vacca et al., 2019). Indeed, we detected changes in the Dp1Tyb hippocampus that can be linked to excitotoxic and neuroinflammatory processes via an increase in the number of microglia and astrocytes as well as the aforementioned significant increases in glutamine and the glutamate/glutamine ratio. The increase in microglia and astrocytes has also been observed in other DS models, such as in Dp(16)1Yey mice, and in humans (Chen et al., 2014; Pinto et al., 2020), highlighting the importance of these cell populations in the correct functioning and development of the brain (Reemst et al., 2016).

We also observed a significant reduction in the number of viable neurons (i.e., NeuN-positive cells) within the hippocampus of Dp1Tyb mice. Similar findings have been reported in humans with DS and in a previous model of DS, the Ts65Dn mouse (Bartesaghi, 2023). However, interpreting results from the Ts65Dn can be challenging, since they harbour up to sixty genes in three copies that are not orthologous to Hsa21 and therefore not directly involved in DS (Duchon et al., 2011). Nevertheless, additional neuronal loss in the brain of individuals with Down syndrome has been associated with Alzheimer-type neuropathology and may, in combination with developmental abnormalities, be associated with accelerated onset of cognitive decline (Wegiel et al., 2022).

Contrary to our expectations, we did not find any significant group differences in synaptic density, as assessed by hippocampal SV2A expression. However, it is worth noting that the existing literature has reported altered synaptic transmission in DS based on other markers, such as dendritic spine counts and morphology (Ferrer and Gullotta, 1990; Suetsugu and Mehraein, 1980) – these studies have also suggested the presence of an altered E/I balance in DS (Jenny A Klein and Haydar, 2022; Souchet et al., 2014). Further analysis, with alternative methods such as functional MRI (fMRI) and using excitatory and inhibitory pre- and postsynaptic markers could shed more light into synaptic alterations in this mouse model of DS.

Finally, we did not observe a significant genotype × sex interaction across the various parameters evaluated. This lack of significance may be attributed to the limited sample size. Notably, previous studies have revealed sex-related disparities in skeletal development in Dp1Tyb animals (Thomas et al., 2020). Specifically, male Dp1Tyb mice exhibit osteopenic phenotypes at an earlier stage than females, while both sexes display osteoporotic phenotypes during early adulthood, mirroring observations in humans with Down syndrome (Carfì et al., 2017; Gavris et al., 2014). Given the scarcity of research on the neurobiological differences between males and females with DS (Johnstone and Mobley, 2023), it is important to conduct further investigations that consider the potential impact of sex on the heterogeneity of findings within the DS population (Andrews et al., 2022).

## 5. CONCLUSION

The findings from our comprehensive imaging and spectroscopic investigation of Dp1Tyb brains confirm a robust relationship between gene triplication and cerebral alterations. The use of this highly representative model, closely resembling the human condition, presents a valuable tool to evaluate the effectiveness of emerging treatments aimed at ameliorating the brain-related pathologies observed in DS. Moreover, our study underscores the imperative for more thorough examinations of synaptic density and function, the involvement of glial cells, and sex-specific disparities within the DS brain. Such investigations hold the potential to identify novel therapeutic targets, ultimately enhancing the quality of life for individuals living with DS.

## ETHICS APPROVAL

Animal work was performed in accordance with the United Kingdom Animal (Scientific Procedures) Act 1986, approved by the Ethical Review panel of the Francis Crick Institute and carried out under Project Licences granted by the UK Home Office. Reporting is based on the ARRIVE Guidelines for Reporting Animal Research (Kilkenny et al., 2010).

## CRediT AUTHORSHIP CONTRIBUTION STATEMENT

Maria Elisa Serrano: Investigation, Methodology, Formal analysis, Visualization, Writing – original draft, Writing – review & editing. Eugene Kim: Investigation, Methodology, Formal analysis, Software, Visualization, Writing – review & editing. Bernard Siow: Methodology, Software, Resources, Supervision. Da Ma: Methodology, Formal analysis, Software, Visualization, Writing – review & editing. Loreto Rojo: Investigation. Camilla Simmons: Investigation. Darryl Hayward: Investigation. Dorota Gibbins: Investigation. Nisha Singh: Conceptualization. Andre Strydom: Conceptualization, review and editing. Elizabeth Fisher: Conceptualization, Resources, Writing – review & editing. Victor Tybulewicz: Conceptualization, Funding acquisition, Resources, Project administration, Supervision, Writing – review & editing. Diana Cash: Conceptualization, Funding acquisition, Resources, Project administration, Supervision, Writing – review & editing.

## Abbreviations

CA: cornu ammonis
Cr: creatine
DG: dentate gyrus
DS: Down syndrome
FDR: false discovery rate
FWE: family-wise error
GABA: gamma-aminobutyric acid
Gln: glutamine
Glu: glutamate
GSH: glutathione
Hsa21: human chromosome 21
IF: immunofluorescence
Ins: *myo*-inositol
Lac: lactate
Mmu: mouse chromosome
MRI: magnetic resonance imaging
MRS: ^1^H spectroscopy
NAA: N-acetyl-aspartate
PCh: phosphocholine
PCr: phosphocreatine
ROI: regions of interest
SEM: standard error of the mean
Taur: taurine
WT: wild-type

## ACKNOWLEDGEMENTS

We would like to thank Abi Gartner and Mary Rutherford for their conversations and valuable insights about the DS pathology in humans and the parallelism between preclinical models – clinical research in this disease.

## DATA AVAILABILITY

Dp1Tyb mice are available through the Jackson Laboratory (strain #037183) and the European Mouse Mutant Archive.

Raw MR data are available on OpenNeuro: https://doi.org/10.18112/openneuro.ds004644.v1.0.0.

## Footnotes

1 https://github.com/dama-lab/multi-atlas-segmentation

2 https://github.com/dama-lab/shape_morphological_analysis

