## Supplemental Figures and Tables for "Investigating Brain Alterations in the Dp1Tyb Mouse Model of Down Syndrome"

**SUPPLEMENTARY DATA**

### SUPPLEMENTARY TABLES

**Supplementary Table 1. Details of antibodies used to perform free-floating immunofluorescence.** In grey, antibodies used for GFAP/NeuN IF and in white, those used for Iba1/SV2A IF.

| | Antibody | Supplier | Catalog number | Dilution |
|---|---|---|---|---|
| *Primary antibodies* | *chicken anti-NeuN* | *Synaptic systems* | *#266006* | *1:200* |
| | *rabbit anti-GFAP* | *DAKO* | *#Z-0334* | *1:1000* |
| | *goat anti-Iba1* | *Abcam* | *#ab5076* | *1:1000* |
| | *rabbit anti-SV2A* | *Abcam* | *#ab32942* | *5µg/ml* |
| *Secondary antibodies* | *Alexa Fluor 488 donkey anti-chicken IgY* | *Jackson Immuno* | *#703-545-155* | *1:600* |
| | *Alexa Fluor 568 donkey anti-rabbit* | *Thermo Fisher* | *#A10042* | *1:600* |
| | *Alexa fluor 488 donkey anti-goat* | *Jackson Immuno* | *#705-605-147* | *1:3000* |
| | *Alexa fluor 568 donkey anti-rabbit* | *Thermo Fisher* | *#A10042* | *1:500* |

**Supplementary Table 2. Comparison of in vivo and ex vivo whole brain (WB) and cerebellum (CB; absolute and relative) volume measurements.** Data are presented as mean ± SEM (m, males; f, females). There was a significant effect of genotype in both WB and absolute CB volumes in ex vivo images (bold, $F_{1,18}$ =20.97, p=0.0039**, $F_{1,18}$ =6.058, p=0.0242*, respectively) and no sex x genotype interaction (2 way ANOVA).

| | *In vivo* | | *Ex vivo* | |
|---|---|---|---|---|
| | WT (7m, 6f) | Dp1Tyb (4m, 5f) | WT (8m, 6f) | Dp1Tyb (5m, 3f) |
| WB ($mm^3$) m | 465.4 ± 4.42 | 443.9 ± 7.85 | **508.0 ± 2.6** | **486.4 ± 8.21** ** |
| WB ($mm^3$) f | 462.6 ± 3.07 | 458.8 ± 10.94 | **507.4 ± 4.53** | **490.7 ± 5.90** |
| CB ($mm^3$) m | 36.5 ± 0.38 | 34.9 ± 0.75 | **37.5 ± 0.35** | **34.4 ± 0.80** * |
| CB ($mm^3$) f | 36.3 ± 0.50 | 36.2 ± 1.47 | **36.8 ± 1.02** | **35.7 ± 0.98** |
| CB % of WB m | 7.8 ± 0.04 | 7.9 ± 0.15 | 7.4 ± 0.05 | 7.1 ± 0.09 |
| CB % of WB f | 7.9 ± 0.10 | 7.9 ± 0.18 | 7.3 ± 0.20 | 7.3 ± 0.12 |

**Supplementary Table 3. Regions with significantly different *in vivo* volume in Dp1Tyb mice compared to WT.** Dp1Tyb mice (n = 9) had a significantly different volume in 26 out of 73 examined ROIs, compared to WT (n= 13). These group differences have been evaluated with a two-way ANOVA (genotype × sex), using the false discovery rate (FDR) to correct for multiple comparisons (p < 0.007, Q = 5%). Regions that are larger in Dp1Tyb brains compared to WT are indicated in red.

| *REGION* | *FUNCTION* | *Difference* | *F* | *p* |
|---|---|---|---|---|
| *Orbital cortex* | *Decision making & executive process* | *WT > Dp1Tyb* | $F_{1,18} = 12.77$ | *0.002* |
| *Prelimbic cortex* | | *WT > Dp1Tyb* | $F_{1,18} = 20.14$ | *< 0.001* |
| *Infralimbic cortex* | | *WT > Dp1Tyb* | $F_{1,18} = 30.47$ | *< 0.001* |
| *Retrosplenial cortex* | *Working memory & spatial memory tasks* | *WT > Dp1Tyb* | $F_{1,18} = 32.48$ | *< 0.001* |
| *Dorsal hippocampus* | | *WT > Dp1Tyb* | $F_{1,18} = 27.09$ | *< 0.001* |
| *Motor cortex* | *Processing of sensorial stimuli* | *WT > Dp1Tyb* | $F_{1,18} = 15.09$ | *0.001* |
| *Sensory Cortex* | | *WT > Dp1Tyb* | $F_{1,18} = 13.54$ | *0.002* |
| *Auditory cortex* | | *WT > Dp1Tyb* | $F_{1,18} = 9.09$ | *0.007* |
| *Olfactory cortex* | | *WT > Dp1Tyb* | $F_{1,18} = 17.88$ | *0.001* |
| *Olfactory tracts* | | *WT > Dp1Tyb* | $F_{1,18} = 21.62$ | *< 0.001* |
| *Piriform cortex* | | *WT > Dp1Tyb* | $F_{1,18} = 10.77$ | *0.004* |
| *Claustrum* | | *WT > Dp1Tyb* | $F_{1,18} = 12.33$ | *0.002* |
| *Parietal cortex* | | *WT > Dp1Tyb* | $F_{1,18} = 12.54$ | *0.002* |
| *Habenula* | *Stress & Anxiety* | *WT > Dp1Tyb* | $F_{1,18} = 12.24$ | *0.003* |
| *Dorsal peduncular area* | | *WT > Dp1Tyb* | $F_{1,18} = 16.28$ | *0.001* |
| *Thalamus* | *Emotional response* | *WT > Dp1Tyb* | $F_{1,18} = 11.98$ | *0.003* |
| *Insular cortex* | | *WT > Dp1Tyb* | $F_{1,18} = 17.64$ | *0.001* |
| *Cingulate cortex* | | *WT > Dp1Tyb* | $F_{1,18} = 29.35$ | *< 0.001* |
| *Amygdala* | | *WT > Dp1Tyb* | $F_{1,18} = 10.01$ | *0.005* |
| *Septum* | | *WT < Dp1Tyb* | $F_{1,18} = 9.53$ | *0.006* |
| *Fourth ventricle* | *Cerebrospinal fluid production* | *WT > Dp1Tyb* | $F_{1,18}= 9.25$ | *0.007* |
| *Internal capsule* | *Cortex-brainstem communication* | *WT > Dp1Tyb* | $F_{1,18} = 10.26$ | *0.005* |
| *Pons* | *Regulation of sleep-wake cycle & autonomic functions* | *WT < Dp1Tyb* | $F_{1,18} = 13.34$ | *0.002* |
| *Pontine reticular nucleus* | | *WT < Dp1Tyb* | $F_{1,18} = 15.17$ | *0.001* |
| *Pedunculopontine nucleus* | | *WT < Dp1Tyb* | $F_{1,18} = 25.56$ | *< 0.001* |
| *Flocculus* | *Motor control – vestibulo-ocular reflex system* | *WT > Dp1Tyb* | $F_{1,18} = 24.79$ | *< 0.001* |

**Supplementary Table 4. Metabolites measured by MRS (mM, except Gln/Glu; mean ± sem).** Data were analysed by two-way (genotype × sex) ANOVA and p values show the effect of genotype, * p<0.05, ** p<0.01, ***p<0.001. Three metabolites depicted by # (glutamine, taurine and glutamine/glutamate ratio) remained significant after correction for multiple comparisons (q-value set as 0.05) controlling the false discovery rate (FDR).

| *MRS metabolites* | **WT** | **Dp1Tyb** |
|---|---|---|
| | males / females | males / females |
| creatine (Cr) | 3.19±0.14 / 2.97±0.10 | 2.92±0.28 / 2.82±0.24 |
| gamma-aminobutyric acid (GABA) | 1.85±0.09 / 1.94±0.08 | 2.00±0.09 / 1.88±0.06 |
| glutamine (Gln) | 1.98±0.05 / 2.02±0.06 | ↑ 3.39±0.88 / 3.31±0.30 ***# |
| glutamine/glutamate (Gln/Glu) | 0.31±0.01 / 0.32±0.01 | ↑ 0.60±0.17 / 0.56±0.07 ***# |
| glutamate (Glu) | 6.30±0.07 / 6.32±0.13 | ↓ 5.88±0.26 / 5.97±0.22 * |
| glutathione (GSH) | 1.39±0.02 / 1.43±0.04 | ↓ 1.38±0.03 / 1.30±0.04 * |
| myo-inositol (Ins) | 3.63±0.11 / 3.91±0.10 | 3.88±0.55 / 3.46±0.31 |
| lactate (Lac) | 1.38±0.06 / 1.34±0.16 | ↑ 1.80±0.35 / 2.01±0.18 ** |
| N-acetyl-aspartate (NAA) | 5.41±0.06 / 5.44±0.06 | ↓ 5.25±0.11 / 5.17±0.12 * |
| phosphocholine (PCh) | 0.75±0.02 / 0.76±0.05 | 0.81±0.10 / 0.80±0.08 |
| taurine (Taur) | 6.91±0.20 / 6.19±0.17 | ↓ 5.33±0.31 / 5.21+0.40 ***# |

**Supplementary Table 5. *Ex-vivo* differences in cerebellar thickness and/or volume between 3-month-old WT and Dp1Tyb mice.** Cerebellar ROIs where we can observe differences in cerebellar thickness and/or volume between WT (n= 13) and Dp1Tyb (n = 9) calculated with a two-way ANOVA (genotype × sex). In red, the statistically significant differences found after multiple comparisons correction using the false discovery rate (FDR) (p < 0.0032, Q = 5%). Abbreviations: lobules of the cerebellar vermis: 3Cb (lobule 3), 4/5Cb (lobule 4/5), 7Cb (lobule 7), 8Cb (lobule 8), 9Cb (lobule 9), 10Cb (lobule 10); lobules of cerebellar hemispheres: Crus 1 (Crus 1 of the ansiform lobule), Crus 2 (Crus 2 of the ansiform lobule), PM (paramedian lobule), Cop (Copula of the pyramis), PFl (Paraflocculus), Fl (Flocculus).

| *REGION* | *MOLECULAR layer* | | *GRANULAR layer* | |
|---|---|---|---|---|
| | *THICKNESS* | *VOLUME* | *THICKNESS* | *VOLUME* |
| *3Cb* | $p < 0.05$ | $F_{1,18} = 34.66$, $p < 0.0001$ | $F_{1,18} = 8.95$, $p = 0.008$ | $F_{1,18} = 18.21$, $p = 0.0005$ |
| *4/5Cb* | $p < 0.05$ | $p < 0.05$ | $p < 0.05$ | $F_{1,18} = 5.08$, $p = 0.037$ |
| *7Cb* | $p < 0.05$ | $F_{1,18} = 7.32$, $p = 0.015$ | $p < 0.05$ | $p < 0.05$ |
| *8Cb* | $F_{1,18} = 10.74$, $p = 0.004$ | $F_{1,18} = 21.84$, $p = 0.0002$ | $F_{1,18} = 10.39$, $p = 0.005$ | $F_{1,18} = 16.69$, $p = 0.0007$ |
| *9Cb* | $p < 0.05$ | $F_{1,18} = 5.87$, $p = 0.026$ | $p < 0.05$ | $F_{1,18} = 17.17$, $p = 0.0006$ |
| *10Cb* | $F_{1,18} = 8.41$, $p = 0.01$ | $F_{1,18} = 8.01$, $p = 0.011$ | $F_{1,18} = 9.65$, $p = 0.006$ | $F_{1,18} = 15.53$, $p = 0.001$ |
| *Crus1* | $p < 0.05$ | $p < 0.05$ | $F_{1,18} = 10.41$, $p = 0.005$ | $F_{1,18} = 7.55$, $p = 0.013$ |
| *Crus2* | $F_{1,18} = 5.32$, $p = 0.033$ | $F_{1,18} = 6.18$, $p = 0.023$ | $p < 0.05$ | $F_{1,18} = 6.44$, $p = 0.021$ |
| *Cop* | $p < 0.05$ | $p < 0.05$ | $p < 0.05$ | $F_{1,18} = 9.24$, $p = 0.007$ |
| *PFl* | $p < 0.05$ | $F_{1,18} = 8.62$, $p = 0.008$ | $p < 0.05$ | $F_{1,18} = 16.64$, $p = 0.0007$ |
| *Fl* | $p < 0.05$ | $F_{1,18} = 14.84$, $p = 0.001$ | $F_{1,18} = 10.54$, $p = 0.005$ | $F_{1,18} = 13.61$, $p = 0.002$ |
| *ALL* | $p < 0.05$ | $F_{1,18} = 7.43$, $p = 0.014$ | $F_{1,18} = 7.18$, $p = 0.015$ | $F_{1,18} = 11.74$, $p = 0.003$ |

*SUPPLEMENTARY FIGURES*

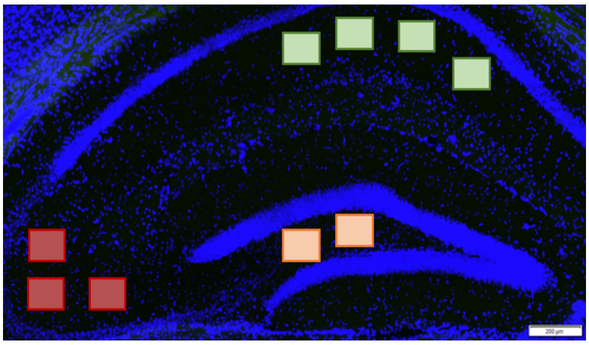

**Supplementary Figure 1. Hippocampal subregions selected for the immunofluorescence (IF) analysis.** Representative hippocampal image after DAPI counterstaining of nuclei and the three ROIs used for the IF analysis: CA1 (green), CA3 (red), DG (orange). Image captured at 10x magnification with a Virtual Slide Microscope VS120 (Olympus Life Science). Scale bar: 200 µm.

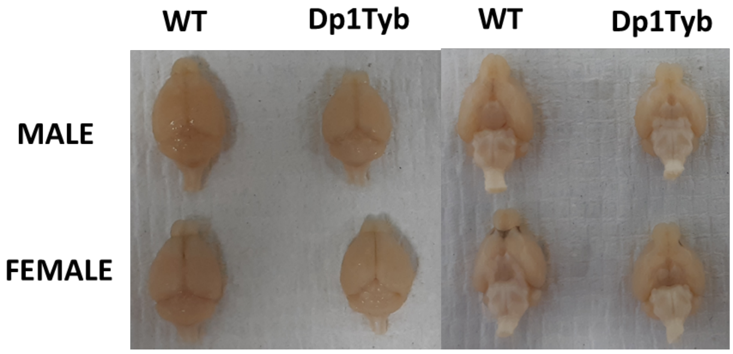


**Supplementary Figure 2. Representative brains of 3-month-old WT and Dp1Tyb mice.** Animals were perfused with heparinized saline and 5% PFA. Photos were taken after brain extraction, showing differences in the shape of representative WT and Dp1Tyb brains, of both sexes. The Dp1Tyb brains are smaller and rounder than the WT ones.

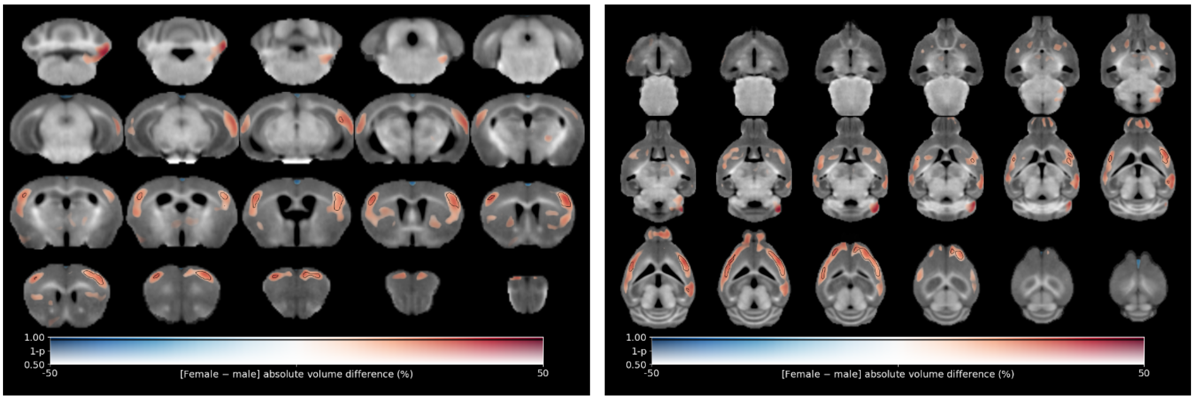


**Supplementary Figure 3. Voxel-wise differences in volume between 3-month-old male and female mice (both genotypes combined together).** Map of voxel-wise differences, derived from in vivo MR images and overlaid on the T1-weighted study-specific template. The map is displayed in the coronal plane (left image, caudal-rostral) and the horizontal plane (right image, ventral-dorsal). The colour of the overlay indicates the percent volume difference (hot colours indicate increased volume in female compared to male), and the opacity of the overlay indicates the significance of the volume difference (regions where the FWE-corrected $p > 0.5$ are completely transparent, and regions where the FWE-corrected $p = 0$ completely opaque). Clusters where the FWE-corrected $p < 0.05$ are contoured in black.

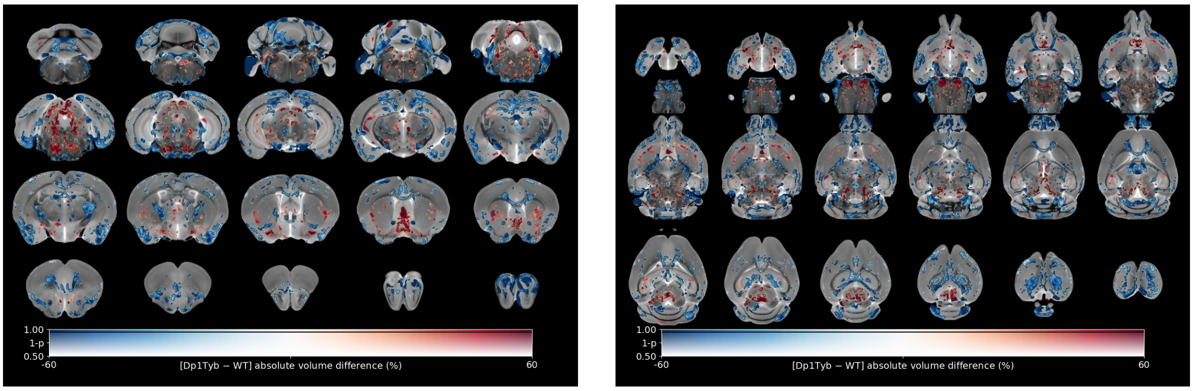


**Supplementary Figure 4. Ex-vivo differences in volume between 3-month-old WT and Dp1Tyb mice.** Map of voxel-wise differences in volume, calculated from ex vivo MR images and overlaid on the Allen mouse brain template. The map is displayed in the coronal plane (left image, caudal-rostral) and the horizontal plane (right image, ventral-dorsal). The colour of the overlay indicates the percent volume difference (cool colours indicate reduced volume in Dp1Tyb mice, n = 9 compared to WT, n = 13), and the opacity of the overlay indicates the significance of the volume difference (regions where the FWE-corrected $p > 0.5$ are completely transparent, and regions where the FWE-corrected $p = 0$ completely opaque). Clusters where the FWE-corrected $p < 0.05$ are contoured in black.
